# Cross-task, explainable, and real-time decoding of human emotion states by integrating grey and white matter intracranial neural activity

**DOI:** 10.1101/2025.11.12.687932

**Authors:** Yuxiao Yang, Wenjun Chen, Yuyang Chen, Ling Ding, Chi Zhang, Hongjie Jiang, Zhoule Zhu, Xinxia Guo, Shuang Wang, Gang Pan, Ning Wei, Shaohua Hu, Junming Zhu, Yueming Wang

## Abstract

Decoding human emotion states from intracranial neural activity is key in developing brain-computer interfaces and new therapies for affective disorders. However, real-world application of decoding requires high performance that integrates neural activity from both grey and white matter, stable generalization across different contexts, sufficient neural encoding explainability, and robust real-time implementation, all of which remain elusive. Here, we simultaneously recorded intracranial electroencephalogram (iEEG) and abundant self-rated valence and arousal scores across two emotion-eliciting tasks in eighteen subjects, forming the largest intracranial neural dataset for emotion decoding. We then developed personalized decoding models within a hybrid deep learning framework involving both self-supervised and supervised components. The models achieved high-performance decoding of continuous valence/arousal states, doubling the R-squared performance of prior EEG/iEEG decoding. Critically, the models significantly improved performance by integrating grey and white matter signals and demonstrated cross-task generalization. The models further revealed shared and preferred mesolimbic-thalamo-cortical subnetworks encoding valence and arousal, showing neurophysiological explainability. Finally, the models realized robust real-time decoding in four new subjects. Our results have implications for advancing emotion decoding neurotechnology toward deployable affective brain–computer interfaces and closed-loop therapeutic systems for affective disorders.

## Main

Emotion represents a fundamental yet complex biological state underlying human survival, social interactions, and mental health^1,2^. To date, there is no consensus on a universal model for emotion^1,2^. Discrete categorical models are adopted by the basic emotion theory^3,4^ while continuous dimensional models are preferred by the cognitive^5,6^ and constructed^7,8^ emotion theory, where valence (a pleasure–displeasure continuum) and arousal (an activation–inactivation continuum) are the most widely recognized core dimension^1,2^. A recent cross-species emotion primitive theory^9,10^ views emotions as evolutionarily conserved central brain states arising from external and internal inputs, being encoded by widespread neural circuits, and causally affecting various output behavioural variables. The central brain state is described by a collection of dimensional features termed emotion primitives, including valence and arousal, that distinguish emotion from other behavioural states or reflexes. The theory describes clear causal links among emotion primitives, neural activity, and behaviour, offering a suitable computational framework for decoding (i.e., statistically inferring) human emotion states from observed neural signals^11–14^, which is crucial for advancing affective brain-computer interfaces^12^, embodied AI^15^, and neuromodulation therapies for affective disorders^16^.

Experimental design, signal modality, and data amount are basic considerations for decoding^11–14,17^. Task-based experiments employing standardized stimuli enable more consistent elicitation and more frequent measurement (usually via self-reports) of emotion than spontaneous reports or clinical interviews under open environments^18^. Numerous task-based neuroimaging studies have investigated emotion encoding circuits^19^, but their low temporal resolution is not suitable for continuous decoding over time^20^. Scalp electroencephalography (EEG) enabled decoding over time but its low spatial and noise resolutions restrict most applications within classifying limited emotion categories^21–23^; only few recent studies explored regression of valence^24–26^. By contrast, intracranial EEG (iEEG) strikes a balance among temporal, spatial, and noise resolutions and has led to more detailed delineation of emotion encoding neural dynamics^27–36^, more accurate emotion classification^37–39^, and fine-grained regression of mood or depression symptom scores^40–59^. While related, mood or symptom is not identical with emotion^7,8,60^ and suffers from sparse data collection (<40 reports per subject, see Discussion), hindering robust decoding evaluation and interpretation. Thus, decoding emotion from task iEEG can provide multiple technological advantages but asks for high performance that integrates useful neural signals, stable generalization across different tasks, sufficient neural encoding explainability, and robust real-time implementation, all of which remain elusive.

First, high-performance decoding requires a computational model integrating useful neural signals. Existing iEEG studies exclusively use grey matter signals for decoding^35–59^ because signals recorded from iEEG channels located in the white matter (referred to as ‘white matter signals’ thereafter for short, see Discussion for detailed clarification) are much weaker, traditionally considered as noise, and disregarded^61^. However, white matter tracts provide the structural basis for inter-regional communication in emotion processing^62,63^. Thus, we hypothesize that integrating grey and white matter iEEG signals can boost decoding. But such integration will need the design of specialized decoding models, especially given the higher neural feature dimensionality resulted from adding white matter signals. Existing iEEG decoding uses linear^41–59^ or classical machine learning models^37–40^, which are likely insufficient to deal with the high dimensionality and nonlinearity of neural dynamics, asking for new decoding models designed from a more advanced deep learning perspective^17^.

Second, generalization across different emotion-eliciting tasks is a critical requirement for decoding. The emotion primitive theory proposes that different stimuli activate a common central brain state that causally leads to downstream responses. This is a core feature of emotion because it allows the individual to quickly sense, process, and respond to a wide range of stimuli across various contexts^9,10^. The theory thus predicts that the neural representations and decoding model associated with emotion primitives, as identified in a single task, should be transferable to apply for other tasks. However, existing decoding studies have exclusively conducted single-task analyses^11–14,17,21–23^, which did not reflect the central role of emotion across diverse contexts. For practical applications, cross-task transferable decoding is necessary for rapid plug-and-play implementation of a trained decoding model across various scenarios^23,64^.

Third, real-world utility of decoding needs the endorsement of a sufficient level of neural encoding explainability provided by the decoding model. Spatially, EEG decoding models have only investigated cortex lobe encoding^21–23^ due to their low resolution; iEEG decoding models have offered more precise, region-level explainability (especially at subcortical regions), but have only pointed to the general importance of mesolimbic regions for encoding composite score representations of mood/symptom^40–59^. A more detailed level of neural encoding explainability should move forward to reveal how different subcortical and cortical regions preferentially encode different emotion primitives, but has remained elusive.

Fourth, real-time validation is the ultimate test of any decoding model if it is to be used in real-world applications such as affective robotics^65^, neurofeedback training^66^, and adaptive neuromodulation^67^. For example, in adaptive deep brain stimulation (aDBS) for affective disorders, emotion-related states need to be decoded from neural activity in real time to provide feedback for delivering optimized stimulation^68–71^.

Unfortunately, existing decoding studies largely remain limited to offline analyses^35–58^, relying on a non-causal cross-validation procedure for decoding evaluation, meaning training data can happen after test data in time; thus, it is difficult to infer if these models are robust enough for real-time applications requiring causal decoding. Also, technical difficulties such as system communication instability, environmental disturbances, and real-time computation constraints have hindered the development of online emotion decoding systems.

Here, we close the above gaps by demonstrating an integrated framework of high-performance, cross-task, explainable, and real-time decoding of emotion states using iEEG in subjects with epilepsy. We first collected wide-coverage iEEG and abundant emotion self-reports (>170 reports per subject) from eighteen subjects participating in two emotion-eliciting tasks (Fig. 1a-c), forming by far the largest human intracranial neural dataset for emotion decoding (Table 1). We then developed personalized decoding models for each individual subject within a hybrid deep learning framework involving both self-supervised and supervised components. The models achieved fine-grained decoding of continuous valence and arousal values in each subject, whose performance was boosted by integrating grey and white matter signals (Fig. 1d). The model achieved a subject-averaged cross-validated R-squared performance of more than twofold compared to prior EEG and iEEG emotion encoding and decoding models (Table 1). We next showed interchangeable and transferable decoding across the two tasks (Fig. 1e). We further explained the decoding models by using them to identify shared and preferred mesolimbic-thalamo-cortical subnetworks encoding valence and arousal (Fig. 1f). We finally implemented the models online and demonstrated robust real-time decoding in four new subjects (Fig. 1g). Overall, our results have implications for advancing real-time and deployable emotion decoding neurotechnology.

**Table 1.** Data amount and decoding performance comparison with state-of-the-art EEG/iEEG-based continuous emotion state decoding. The underline represents state-of-the-art value.

| Signal | Year | Previous works | Subjects' clinical conditions | # of subjects | Contacts used for decoding per subject | Total # of contacts (total coverage) | # of labels per subject | Total # of labels (sample size) | Regression CC (cross-validated?) | CC difference from existing average | Regression R<sup>2</sup> (cross-validated?) | R<sup>2</sup> difference from existing average |
| --- | --- | --- | --- | --- | --- | --- | --- | --- | --- | --- | --- | --- |
| EEG | 2022 | Zhang et al.<sup>24</sup> | Healthy | 24 | 32 | 768 | 10 | 239 | 0.47 | 0.01↑ | 0.22 (Yes) | 0.01↑ |
|  | 2023 | Ding et al.<sup>25</sup> | Healthy | 24 | 32 | 768 | 10 | 239 | 0.47 | 0.01↑ | 0.22 (Yes) | 0.01↑ |
|  | 2024 | Ding et al.<sup>26</sup> | Healthy | 24 | 32 | 768 | 10 | 239 | 0.51 | 0.05↑ | 0.26 (Yes) | 0.05↑ |
| iEEG | 2016 | Huebl et al.<sup>35</sup> | Depression | 8 | 8 | 64 | 3 | 24 | 0.61 | 0.15↑ | 0.37 (No) | 0.16↑ |
|  | 2023 | Sonkusare et al.<sup>36</sup> | Epilepsy | <u>27</u> | 4 | 108 | 15 | 405 | 0.36 | -0.10↓ | 0.13 (No) | -0.08↓ |
| **Meta-analysis average<sup>133</sup> of previous studies** |  |  |  | 21 | 23 | 495 | 11 | 229 | 0.46 | 0.00 | 0.21 | 0.00 |
| **Ours** |  |  | **Epilepsy** | **18** | **<u>120</u>** | **<u>2160</u>** | **<u>47</u>** | **<u>845</u>** | **<u>0.70</u>** | **<u>0.24</u>**↑ | **<u>0.49</u>** (Yes) | **<u>0.28</u>**↑ |

**Fig. 1.**
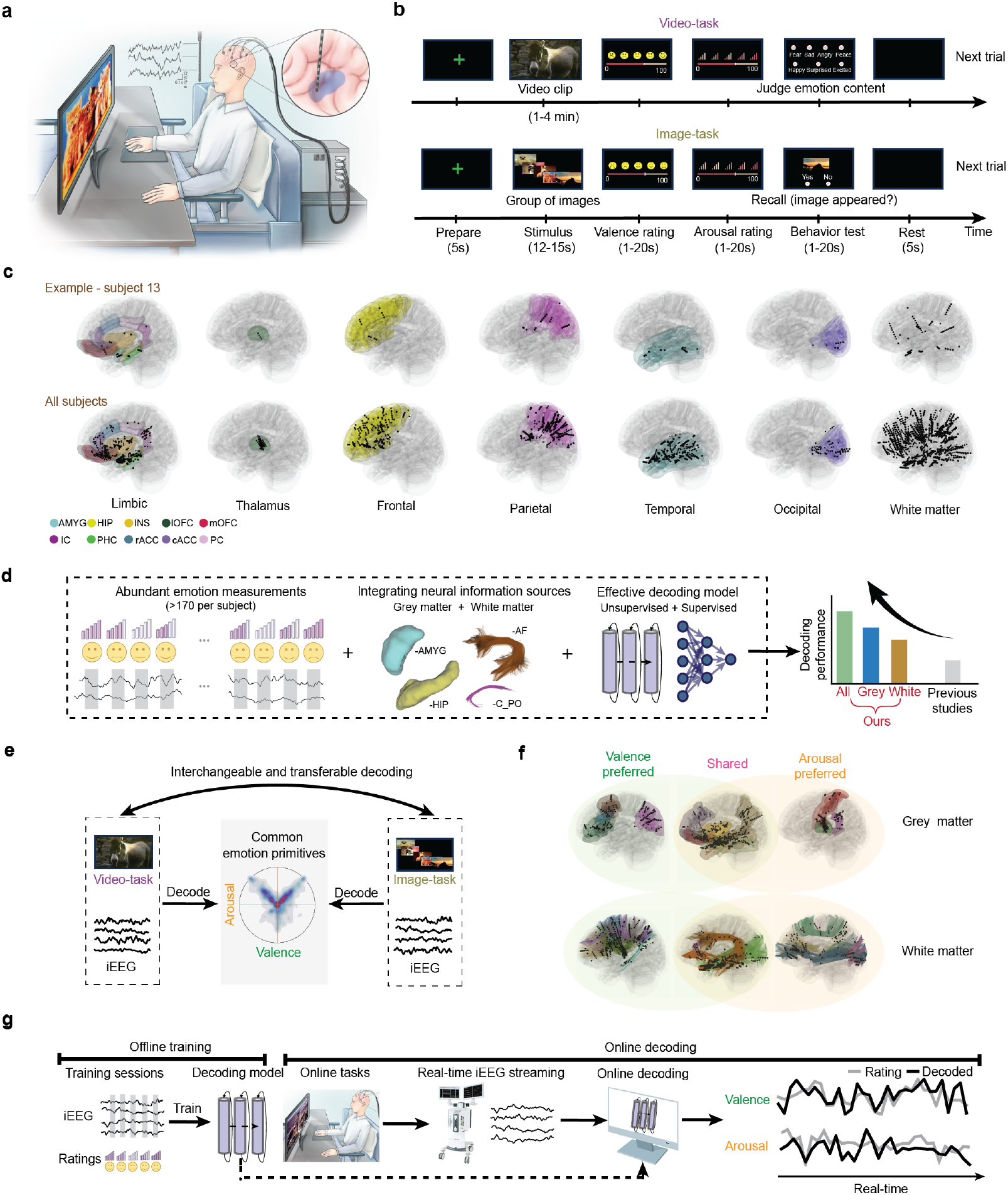
Overview of experimental design, neural recording, and decoding analyses. **a**, Experimental setup for emotion elicitation and iEEG recording. Subjects are instructed to view image or video stimuli and self-report valence and arousal ratings. iEEG signals are continuously recorded. **b**, Schematic diagrams of example trials of the video-task and image-task (see Methods for details). **c**, Top, iEEG contact locations in a representative subject. Bottom, summary of all iEEG contact locations across the population. Each black dot represents a grey matter recording contact overlaid on the DKT and ASEG atlases with different colors representing different grey matter regions. Each black dot without color overlay represents a white matter recording contact. Region name abbreviations: hippocampus (HIP), insula (INS), amygdala (AMYG), lateral orbitofrontal (lOFC), medial orbitofrontal (mOFC), caudal anterior cingulate (cACC), rostral anterior cingulate (rACC), isthmus cingulate (IC), posterior cingulate (PC), parahippocampal (PHC). **d-g**, Summary of the main analyses and findings. **d**, We collected abundant emotion measurements as self-report valence and arousal ratings, took both the grey matter and white matter signals as useful neural information sources, and designed specialized deep learning models to integrate such information, together leading to high-performance decoding. **e**, Our model enabled interchangeable and transferable decoding of the common neural representations of emotion primitives across the two tasks. **f**, Our decoding models achieved a reasonable level of encoding explainability, revealing shared and preferred mesolimbic-thalamo-cortical subnetworks encoding valence and arousal. **g**, New subjects were instructed to perform image-tasks and video-tasks online while iEEG data were streamed to a fixed, offline-trained decoding model for real-time prediction of valence and arousal states.

## Results

### Overview of experimental design, neural recording, and decoding model

We recruited 18 subjects with epilepsy undergoing iEEG monitoring for seizure localization (Fig. 1a and Supp. Table 1). Across 2.27±0.24 days, each subject completed 4.72±0.03 (mean±s.e.m.) rounds of experiments, each comprising three interleaved sessions of two types of emotion-eliciting tasks—static image-viewing task (image-task) and dynamic video-viewing task (video-task) (Fig. 1b, see Methods, Supp. Fig.1, and Supp. Table 2). Each session consisted of 20-23 image-task trials or 7-10 video-task trials, where the subject self-rated valence and arousal states on visual-analog-scales (VAS) spanning 0-100 for each trial. Subjects not fully engaged were removed from further analyses (see Methods), resulting in 18 and 14 subjects included in the offline video-task and image-task decoding analyses, respectively (Supp. Fig. 2 and Supp. Table 3). Overall, after trial inclusion validation, for each subject, 124±7.25 (valence) and 132.36±6.85 (arousal) trials of image-task and 46.94±1.63 (valence) and 47.17±1.52 (arousal) trials of video-task were included in subsequent decoding (see Methods), where the self-rated scores spanned reasonable ranges (Supp. Fig. 3, Supp. Tables 4 and 5). The subjects’ rated scores showed a typical ‘boomerang’ shape in the valence-arousal space, consistent with prior studies^72^ (Supp. Fig. 3a). Another 4 subjects with epilepsy were recruited in the online experiment (see Methods). iEEG signals were recorded concurrently with the tasks (Fig. 1c, see Methods). Across subjects, iEEG contacts were implanted in mesolimbic, subcortical, widespread cortical regions, and white matter tracts, covering 41 grey matter regions based on the Desikan-Killiany-Tourville (DKT)^73^ and FreeSurfer ASEG^74^ atlases and 26 white matter tracts based on the Human-Connectome-Project (HCP)^75^ atlas (Supp. Figs. 4 and 5, Supp. Tables 6, 7 and 8). Compared with existing EEG and iEEG emotion decoding studies, our dataset has the widest electrode coverage, largest number of self-rated emotion labels per subject, and largest total sample size (Table 1).

Our dataset of widespread iEEG signals and abundant emotion measurements allowed in-depth decoding and explainability analyses, as expanded in the next few sections. The foundation for these analyses is the construction of decoding models that map neural activity to valence and arousal ratings. The paradox between the high-dimensional spatial-spectral-temporal neural features and the relatively sparse ratings (although they were much denser than existing studies) makes direct supervised learning difficult, but the large amount of unlabeled iEEG data enables sufficient self-supervised learning. We thus designed a hybrid deep learning model with both self-supervised and supervised components for dimensionality reduction and decoding (Fig. 2a, see Methods). First, we extracted iEEG spectral powers at *δ* (1-4 Hz), θ (4-8 Hz), α (8-13 Hz), β (13-30 Hz), low γ (30-70 Hz), and high γ (70-200 Hz) bands every 1 second for each channel to form a neural feature vector time-series per subject (feature dimension of 708±35.89; total time step of 3710.71±217.37 and 3965.07±209.55 in image-task and 5748.28±240.63 and 5781.00±235.58 in video-task for valence and arousal, respectively). Second, we used a supervised F-statistic-driven method to identify the top discriminative features. Third, we used a self-supervised long short-term memory (LSTM) autoencoder to capture nonlinear neural dynamics through low-dimensional representations that enable one-step-ahead prediction of future features. We chose LSTM over other models, such as the Transformer, because it delivered comparable prediction performance with a simpler architecture (Supp. Fig. 7). Fourth, we built supervised multi-layer perceptron (MLP) modules to use these low-dimensional representations to predict (via regression) fine-grained continuous valence/arousal ratings or categorize (via classification) coarse-grained discretized valence/arousal states. Implementation details are provided in Methods and Supp. Figs. 6 and 7.

**Fig. 2.**
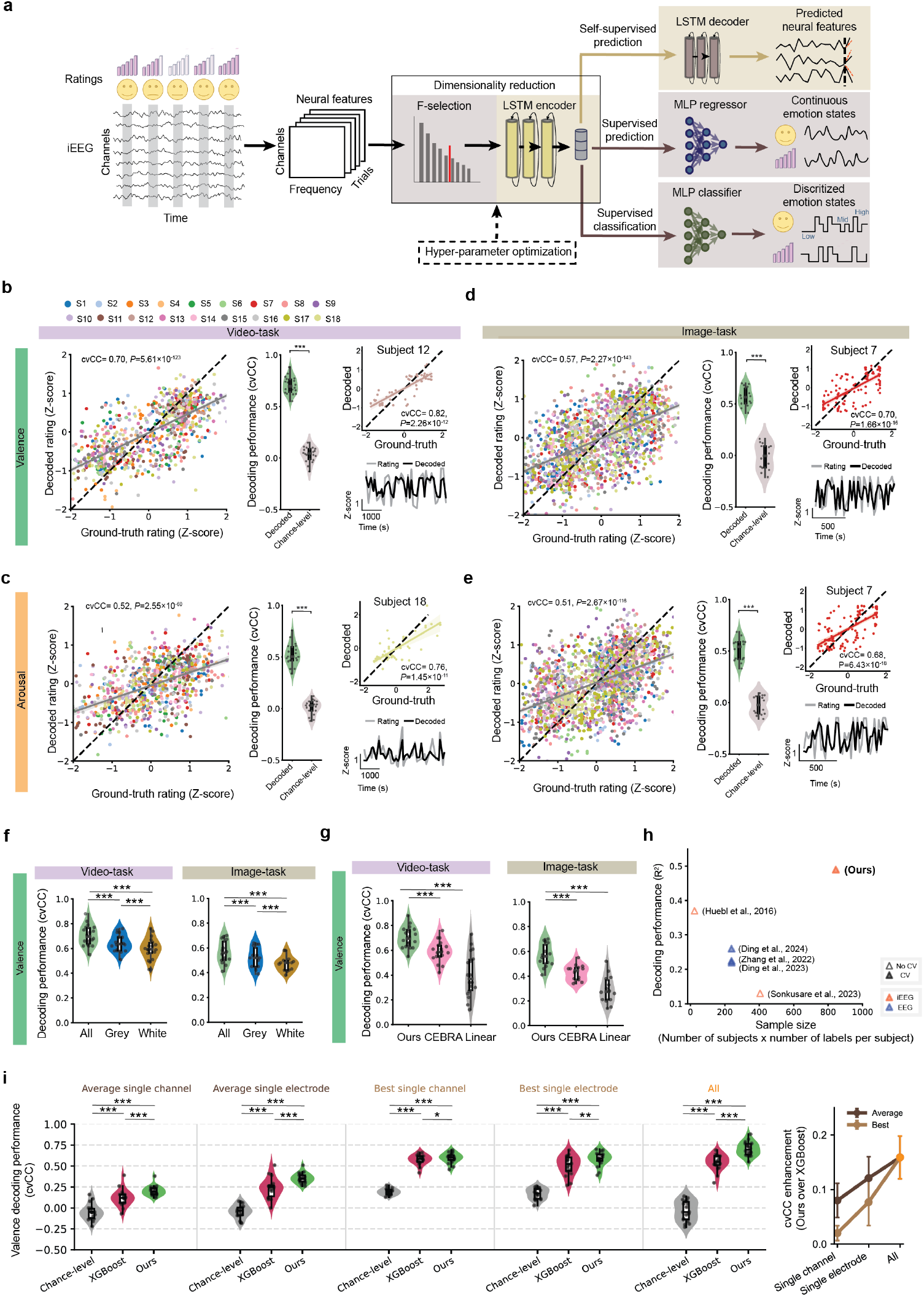
High-performance single-task decoding of valence and arousal states by personalized decoding models designed within a hybrid deep learning framework involving both self-supervised and supervised components. **a**, The data processing pipeline and structure of the neural decoding model, which mainly consists of dimensionality reduction by a self-supervised LSTM autoencoder and supervised learning by an MLP regressor/classifier. **b**, Cross-validated valence decoding results in the video-task. Left, the population-pooled cross-validated regression decoding results, where distinct colors represent different subjects (the colors remain consistent for all following analyses), each dot represents the decoded rating of a single trial (*n*=845), the grey line represents linear fitting of all decoding results with the shaded area representing the 95% confidence interval, and the dotted black line indicates the 45-degree reference line of perfect decoding. Same plotting format holds for subsequent linear fitting. Note that to pool the results across subjects, the ground-truth ratings (horizontal axis) and decoded ratings (vertical axis) are both z-scored by the distribution of the ground-truth ratings within each subject. Middle, the pooled decoding performance significantly outperformed chance-level (*N*=18). In each violin plot, each dot represents a subject. The width of the shape represents the data distribution density. The inner central line represents the median, the box spans the interquartile range (IQR), and whiskers extend to 1.5×IQR. n.s.: *P*>0.05; *: *P*<0.05; **: *P*<0.005; ***: *P*<0.0005. Subsequent violin plots follow the same format. Right top, example decoding results of a high-performance subject (see Supp. Fig. 11 for other subjects). Right bottom, the corresponding temporal traces of decoded ratings and ground-truth ratings for the example subject. **c**, Same as **b** but for cross-validated arousal decoding results in the video-task. **d-e**, Same as **b-c** but for the image-task (*N*=14, *n*=1736). **f**, Population-level valence decoding performance comparison in the video-task and image-task of (1) using all iEEG contacts; (2) using only iEEG contacts in grey matter; (3) using only iEEG contacts in white matter (see Supp. Fig. 14 for arousal results). **g**, Valence decoding performance comparison of our model with the deep learning-based CEBRA decoding model and the linear control model (see Supp. Fig. 12 for arousal results). **h**, Comparison of our decoding performance with prior iEEG and EEG studies investigating continuous emotion state decoding. Each triangle represents the average performance of an individual study: hollow triangles denote studies without cross-validation, and solid triangles denote studies with cross-validation, color-coded by signal modality. **i**, Comparison of our decoding model with the XGBoost decoding model. Left, comparisons under five conditions: (1) the average performance of using a single channel; (2) the average performance of using a single electrode; (3) the best performance among single channels; (4) the best performance among single electrodes; (5) the performance of using all channels. Right, cvCC improvement of our model over XGBoost when moving from a smaller number of iEEG channel to larger numbers of channels. The dot represents the mean with whiskers representing the 95% confidence interval. Subsequent error bar plots follow the same format.

We trained, validated, and tested the above decoding model separately for each subject using rigorous 10-fold leave-trials-out cross-validation (Supp. Fig. 6; including cross-validating the model’s hyperparameters, see Methods), leading to a personalized decoding framework. The primary decoding performance measures were the cross-validated correlation coefficient (cvCC) and the cross-validated R-squared (cvR^2^) in predicting the continuous valence/arousal ratings and the secondary measure was the cross-validated classification accuracy (cvACC) of high, median, and low valence/arousal states. Other typical regression and classification measures were also computed for completeness. Statistical significance of decoding was assessed by applying the identical framework to randomly permuted valence/arousal ratings, with permutation *P*-values quantifying the probability of random decoding performance (chance-level) exceeding the true decoding performance. All *P*-values were corrected with false discovery rate (FDR) control wherever needed.

### High-performance single-task decoding of emotion states by integrating grey and white matter signals

Our personalized models enabled high-performance decoding of valence and arousal states in distinct emotion-eliciting tasks. For video-task, the personalized model significantly decoded continuous valence ratings in each subject, with high cvCC values ranging from 0.55 to 0.88 and cvR^2^ ranging from 0.29 to 0.69 (*n* = 34 to 56 per subject, where *n* represents the number of ratings; FDR-corrected permutation *P* < 0.008 for each subject; Fig. 2b; Supp. Fig. 8), as confirmed by the strong consistency of temporal decoding traces with the ground-truth rating pattern (Fig. 2b). At the population level, the pooled cvCC was 0.70±0.02, cvR^2^ was 0.49±0.03, significantly larger than chance-level (*N* = 18, where *N* represents the number of subjects, *n* = 845 across subjects, *P* = 7.63×10^−9^; Fig. 2b; Supp. Fig. 9a). For arousal, the individual cvCC ranged from 0.34 to 0.76 and cvR^2^ ranged from 0.10 to 0.57 (FDR-corrected *P* < 0.03 for each subject; Fig. 2c; Supp. Fig. 8) and the population-pooled cvCC was 0.52±0.03, cvR^2^ was 0.26±0.03 (*P* = 7.63×10^−9^; Fig. 2c; Supp. Fig. 9a). Moreover, a simple variant of our model enabled significant classification of high, median, and low valence/arousal states with individual-level cvACC ranging from 0.65 to 0.83 and 0.59 to 0.81, respectively (FDR-corrected *P* < 0.0008 for each subject; Supp. Fig. 10), and pooled cvACC reaching 0.75±0.01 and 0.70±0.01, respectively (*P* < 0.01 for both cases; Supp. Fig. 10). A coarser classification of two-class high and low valence/arousal states led to population-pooled cvACC of 0.81±0.01 and 0.74±0.01, respectively (*P* < 0.01 for both cases; Supp. Fig. 10). Our model outperformed current state-of-the-art EEG and iEEG emotion decoding models, achieving more than double the cvR^2^ (0.49 vs. a meta-analysis average of 0.21), and was validated with a nearly four times larger sample size (*n* = 845 vs. a meta-analysis average *n* = 229) (Fig. 2h and Table 1). Other typical regression and classification performance measures for each subject were included in Supp. Table 9.

Similar results held for image-task decoding. First, our model significantly decoded valence/arousal, with cvCC ranging from 0.41 to 0.70 and 0.33 to 0.68, respectively, for each subject (*n* = 61 to 174 and 69 to 175 per subject, respectively; FDR-corrected *P* < 0.0007 and *P* < 0.0046, respectively; Fig. 2d and 2e; Supp. Fig. 11; Supp. Table 10) and a population-pooled cvCC of 0.57±0.02 and 0.51±0.03, respectively (*N* = 14, *n* = 1736 and 1853 across subjects, respectively; *P* = 1.22×10^−4^ for both; Fig. 2d and 2e, Supp. Fig. 9b). Second, the individual cvACC of three-class valence and arousal classifications ranged from 0.62 to 0.79 and 0.53 to 0.8, respectively (FDR-corrected *P* < 0.006 for each subject; Supp. Fig. 10). The population-pooled cvACC achieved 0.67±0.01 and 0.66±0.02, respectively (*P* < 0.01 for both cases; Supp. Fig. 10; Supp. Table 10).

It should be noted that we separately trained and validated decoding models for valence and arousal states in each subject. Theoretically, valence and arousal are two major but not necessarily independent emotion primitives. As reflected by the self-reported valence and arousal scores across all subjects, they showed a typical “boomerang” pattern, with a linear correlation R-squared of 0.15±0.04 in video-task and 0.15±0.06 in image-task, respectively, and a higher quadratic correlation R-squared of 0.48±0.05 in video-task and 0.44±0.06 in image-task, respectively (Supp. Fig. 3a). In terms of the neural-decoded valence and arousal states, they showed a similar “boomerang” pattern, with a linear correlation fitting R-squared of 0.13±0.04 in video-task and 0.14±0.06 in image-task, respectively, and a quadratic correlation R-squared of 0.40±0.05 in video-task and 0.35±0.07 in image-task, respectively (Supp. Fig. 3b). The results show the behavioral consistency of our decoding methods and suggest a nonlinear correlation between the neural states representing valence and arousal (see Discussion).

The high decoding performance was achieved by integrating grey and white matter signals, outperforming each single type alone. While white matter signals had smaller amplitudes, they were not pure noise but contained substantial information about emotion (Supp. Fig. 14): they decoded valence and arousal significantly better than chance-level across all subjects (*P* < 0.0002 in all cases; Fig. 2f), although the performance was slightly worse than grey matter (valence cvCC, 0.60±0.02 vs. 0.64±0.02 and 0.47±0.02 vs. 0.52±0.02; arousal cvCC= 0.46±0.02 vs. 0.49±0.02 and 0.43±0.02 vs. 0.47±0.03 in video-task and image-task, respectively). However, the decoding performance of all combined signals significantly outperformed either signal type alone (Fig. 2f, *P*<0.0005 in all cases). Moreover, the improvement from adding white matter signals correlated with their structural connectivity pattern to emotion-related grey matter regions (Supp. Note 1 and Supp. Figs. 15-17). These results indicate that white matter signals provide complementary information that augments grey matter signals, boosting the decoding (see Discussion).

In addition, we conducted several control analyses to confirm the decoding effectiveness (Fig. 2g). First, since prior iEEG studies have exclusively used linear dimensionality reduction and regression models (see Discussion), we replaced the MLP module with regularized linear regression and then replaced the LSTM autoencoder module with linear principal component analysis, leading to a decrease of video-task valence cvCC from 0.70±0.02 to 0.52±0.03, and further to 0.40±0.04 (a similar decrease held for image-task and arousal results; Supp. Note 2 and Supp. Fig. 12). This confirms the necessity of nonlinear dimensionality reduction and nonlinear learning for more precise decoding. Second, we compared with the recent deep learning-based CEBRA algorithm^76^ designed for general decoding purposes and found that our model achieved significantly better performance (video-task valence 0.70±0.02 vs. 0.59±0.02, *P* = 7.63×10^−9^; similar results held for image-task and arousal; Supp. Note 2 and Supp. Fig. 12), showing the necessity of explicit temporal neural dynamics prediction in our LSTM encoder. Third, to exclude the influence of epileptic activity^77^, we extracted interictal discharge rates as the features, repeated the same decoding analyses, and found that decoding failed (mean population-pooled cvCC < 0.14, *P* >0.06 in all cases; Supp. Note 3; Supp. Fig. 13). Fourth, to exclude the possibility that the larger number of included features purely contributed to the improvement of combined decoding, we randomly removed iEEG channels in the combined case to match that of grey or white matter decoding alone and still found significantly improved decoding performance (Supp. Note 4 and Supp. Fig. 14b).

Finally, potential clinical applications of intracranial emotion decoding usually rely on implantable devices with a limited number of channels/electrodes and computational power. We thus investigated how our model performed compared with a typical simpler on-chip model, i.e., the eXtreme Gradient Boosting model (XGBoost)^78^, under the ideal scenario of using the full multi-site recordings and the clinical scenarios of single channel decoding (this only requires recording from a focal neural ensemble) or single electrode decoding (this only requires a single implantation trajectory) (Fig. 2i, left). We took the valence decoding of the video-task as the typical scenario since it generally resulted in better performance. In all cases, both our model and XGBoost significantly outperformed the chance-level (*P<0*.*0004* in all cases), confirming the decoding capability of the simpler model and the decoding consistency of our own model. As expected, both our model and XGBoost showed an increasing trend when moving from average single-channel/single-electrode, to best single-channel/single-electrode, and finally to the full multi-site recordings. Notably, in all cases, our model significantly outperformed XGBoost (*P<0*.*02* in all cases), confirming the necessity of our specialized model structure design. The improvement of our model over XGBoost showed an increasing trend when moving from a smaller number of iEEG channels to larger numbers of channels (Fig. 2i, right), where the largest difference was observed in the case of using all channels (Ours: 0.70±0.02 vs. XGBoost: 0.54±0.02, *P* = 0.0003); such a trend suggests the trade-off between decoding performance and model complexity is an important consideration in practical clinical applications (see Discussion).

### Interchangeable, transferable, and temporally-stable decoding across distinct emotion-eliciting tasks

We have demonstrated that emotion states can be effectively decoded for the video-task and image-task. With the assumption that different emotion stimuli activate a common central brain state, we next investigated whether the neural representations of the central brain state enable interchangeable, transferable, and temporally-stable decoding across tasks. The cohort here comprised the 14 subjects who were included in both tasks.

We first investigated whether a model trained by source-task data could be directly interchanged to decode emotion states in the target-task. We trained three models (Supp. Fig. 20): (1) an intra-task model (trained on target-task data), (2) an inter-task model (trained on source-task data), and (3) a random model (trained with randomly permuted target-task labels). We evaluated these models using the same target-task test data with 10-fold leave-trials-out cross-validation (see Methods). Taking image-task as the source-task and video-task as the target-task as an example, the inter-task model achieved a population-level decoding cvCC of 0.34±0.03 for valence (*n* = 643), significantly outperforming the random model (*P*=1.22×10^−4^; Fig. 3a), though the intra-task model, as expected, achieved the best performance. Consistently, t-SNE visualization of the hidden layer embedding of the classification MLP in each subject demonstrated enhanced cluster separation of the inter-task model compared to the random model (Fig. 3b and Supp. Fig. 18) with significantly higher clustering efficacy values (Calinski-Harabasz index, CHI, Fig. 3c), while the intra-task model remained optimal. Similar results held for arousal decoding (Supp. Fig. 19) and for swapping source/target tasks (Supp. Fig. 20). Such interchangeable decoding indicates common neural representations across tasks and the possibility of fine-tuning the inter-task model for better decoding the target-task.

**Fig. 3.**
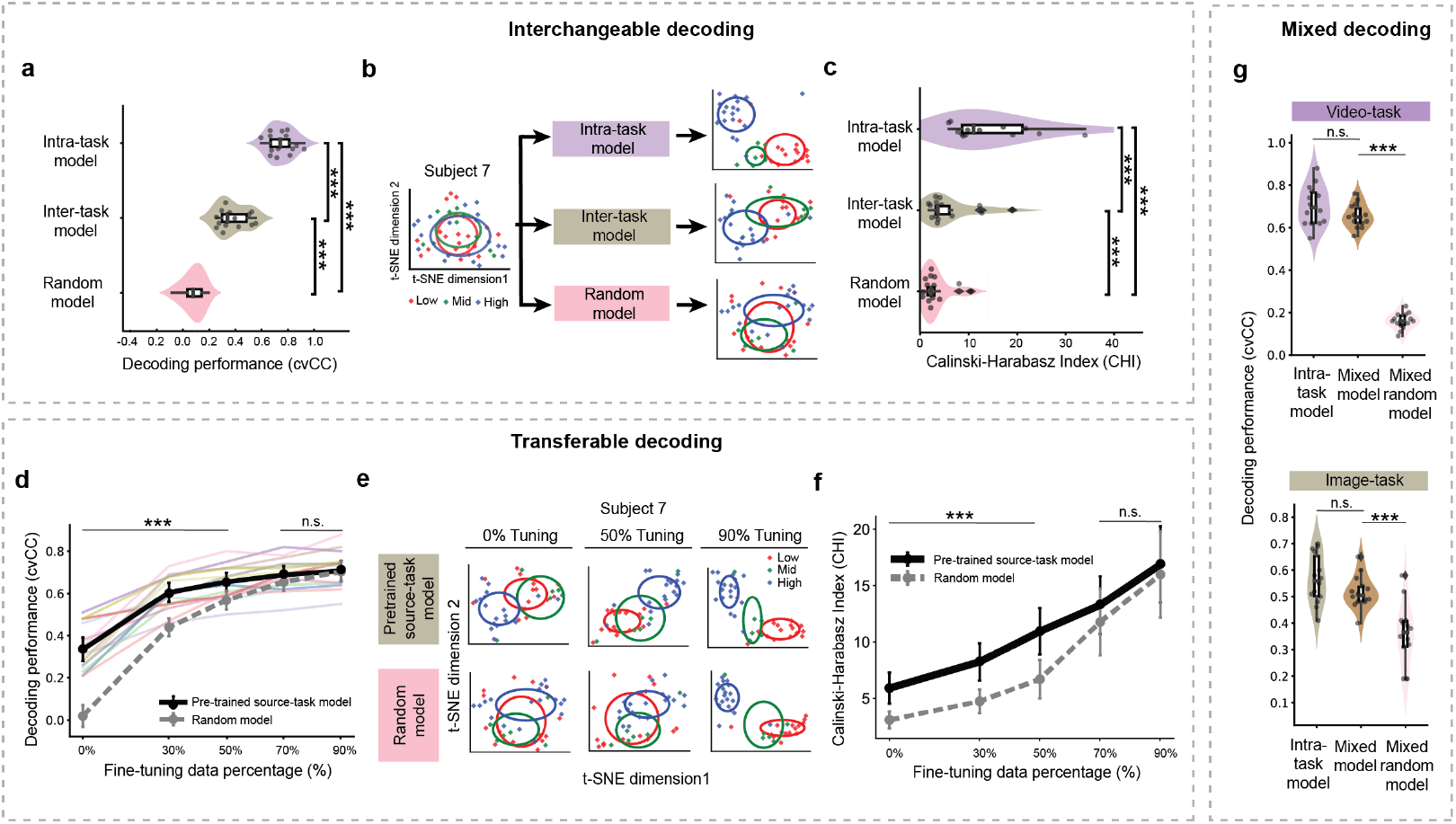
Interchangeable and transferable decoding across distinct emotion-eliciting tasks. **a**, Decoding performance comparison of the intra-task, inter-task and random models across the population. **b**, t-SNE visualization comparison of the three models in **a** for an example subject (see Supp. Fig. 18 for other subjects). Left, 2-dimensional t-SNE visualization of input neural features before decoding, with dots representing neural feature samples, colors representing low, median, and high valence groups, and ellipses representing Gaussian clustering boundaries. Right, t-SNE visualization of the neural embeddings extracted by the three models. **c**, Quantitative comparison of clustering efficacy. Higher CHI values represent better cluster separation. **d**, Decoding performance comparison between fine-tuning the two models with varying proportions of target-task training data. Each colored line represents a single subject, and the solid black line represents the mean with whiskers representing the 95% confidence interval. **e**, 2-dimensional t-SNE visualizations of the fine-tuning process in a single example subject (see Supp. Fig. 21 for other subjects). **f**, CHI clustering efficacy of the fine-tuning process. **g**, Decoding performance comparison between the intra-task model, the mixed model, and mixed random control.

Therefore, we explored whether the inter-task model can serve as a ‘pretrained source-task’ model for rapid fine-tuning by the target-task data within a transfer learning framework (Fig. 3d-f; Supp. Fig. 20). Across the population, when fine-tuned with only 30% of target-task trials, the model achieved a sufficiently good cvCC of 0.63±0.02 for valence, then saturated at 50% of trials with a cvCC of 0.68±0.02, comparable to the intra-task performance of 0.70±0.02 (*N* = 14, *n* = 570, *P* = 0.05; Fig. 3f). The performance surpassed fine-tuning from the random model at each data amount below 50% (FDR-corrected *P* < 0.036; Fig. 3d). Visualization and cluster metrics confirmed the progressive refinement of embeddings (Fig. 3e-f and Supp. Fig. 21). Similar results held for arousal (Supp. Fig. 19) and reversed task roles (Supp. Fig. 20). To confirm the temporal stability of transfer decoding, we additionally performed a causal analysis. The pre-trained source-task model was fine-tuned using the initial round of target-task data (approximately 20% of data). The decoding performance remained high and stable across subsequent rounds (time elapsed from initial round: 15.03±3.03 hours, range: 1–44 hours; FDR-corrected *P* < 0.0042 at each elapsed time in all cases; Supp. Fig. 22), showing the feasibility of rapid and stable transfer.

To further confirm the existence of shared neural representation patterns across different tasks, we conducted a control analysis by mixing video-task and image-task data for training a common mixed decoding model and testing it for separate decoding of video-task and image-task with 10-fold leave-trials-out cross-validation (see Methods; Supp. Fig. 23). We found that the mixed model significantly decoded valence, performing similarly with the intra-task models (*P* > 0.19 in both cases; Fig. 3g); by contrast, mixing video-task (or image-task) and random data for training led to significantly worse decoding (*P* < 2.44×10^−4^ in both cases; Fig. 3g). Similar results held for arousal decoding (Supp. Fig. 23).

### Spatial encoding explanation: Shared and preferred grey matter regions encoding valence and arousal

Having demonstrated effective cross-task decoding, we used the personalized decoding model to explain population-level encoding patterns of valence and arousal. It is worth noting that our primary analytical approach is fitting personalized decoding models for each subject rather than fitting a single decoding model by aggregating data from all subjects (as limited by the subjects’ electrode implantation heterogeneity, see Discussion). Nevertheless, aggregating results across the individually-fitted models can still reveal some common population-level encoding patterns, which constitute our secondary analytical approach detailed below.

First, in each subject, we identified top-decoding grey matter subnetworks from all implanted regions. Previous studies have indicated multi-region subnetworks that reflect emotion and mood. We thus identified subject-specific top-decoding subnetworks of valence (or arousal) by using a greedy algorithm that starts with a single region and gradually adds regions to form best-performing subnetworks with increasing sizes (with a rigorous nested cross-validation procedure to prevent overfitting, see Methods). In each subject, the typical pattern is that as the number of included regions increased, the decoding performance gradually increased and saturated at around four regions, reaching sufficiently good decoding compared with best-possible decoding of using all implanted regions (Supp. Fig. 24). Such observations were confirmed by the average pattern across subjects (four-region cvCC = 0.55±0.02 vs. all-region cvCC = 0.60±0.01, *P* = 0.06; arousal: four-region cvCC = 0.47±0.02 vs. all-region cvCC = 0.52±0.02, *P* = 0.13; Fig. 4a and 4d).

**Fig. 4.**
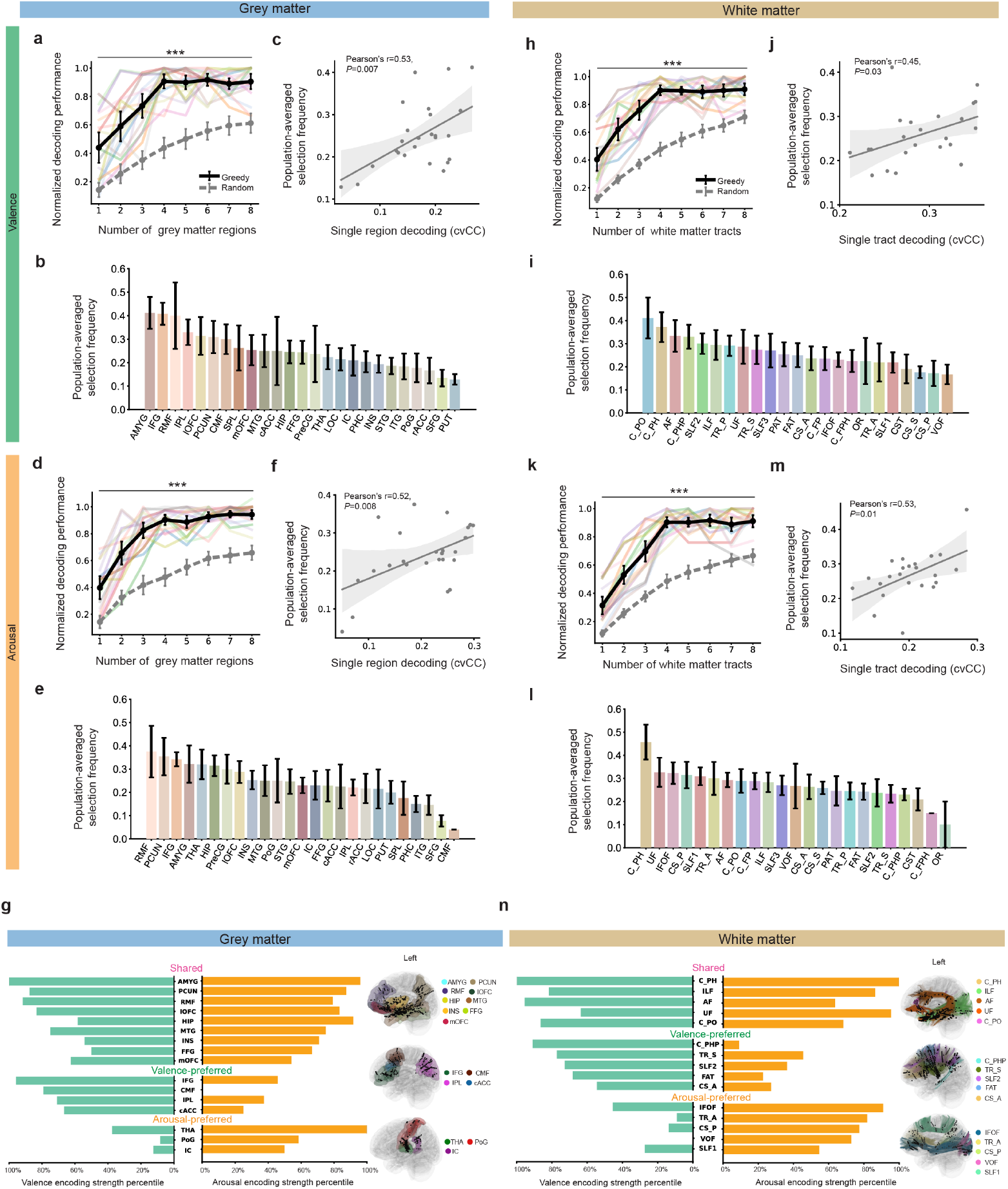
Grey matter regions/white matter tracts differentially encoding valence and arousal. **a**, Valence decoding performance of models trained with an increasing number of grey matter regions as selected by our nested cross-validation procedure and greedy algorithm. Each color represents a single subject where the decoding cvCC is normalized by dividing the cvCC of using all implanted grey matter regions. The solid black line represents population average. The grey line represents chance-level performance of randomly selecting regions for comparison. **b**, Selection frequency of each grey matter region averaged across all subjects. Region name abbreviations: hippocampus (HIP), insula (INS), amygdala (AMYG), lateral orbitofrontal (lOFC), medial orbitofrontal (mOFC), caudal anterior cingulate (cACC), rostral anterior cingulate (rACC), isthmus cingulate (IC), parahippocampal (PHC), superior frontal (SFG), inferior frontal (IFG), caudal middle front (CMF), rostral middle front (RMF), precentral (PreCG), inferior parietal (IPL), superior parietal (SPL), precuneus (PCUN), postcentral (PoG), middle temporal (MTG), superior temporal (STG), inferior temporal (ITG), fusiform (FFG), thalamus (THA), putamen (PUT), lateral occipital (LOC). See Supp. Table 6 for the full list of region name abbreviations. **c**, Pearson correlation between the population-averaged region selection frequency in **b** and the single-region valence decoding performance, with each dot representing a grey matter region and the solid line representing the linear fitting. Population-level key-encoding grey matter regions are defined by combining population-averaged region selection frequency and the single-region decoding performance as detailed in Methods. **d–f**, Captions are the same as **a–c** but for arousal. **g**, Left, division of the population-level key-encoding grey matter regions into the shared group, the valence-preferred group, and the arousal-preferred group. Right, anatomical visualization of the corresponding key-encoding grey matter regions (shaded colors) and electrode implantation locations (black dots) for each group across all subjects. From top to bottom: shared, valence-preferred, and arousal-preferred. **h–n**, panel captions are the same as **a–g** but for key white matter tract distributions. White matter tract name abbreviations: frontal parahippocampal cingulum (C_FPH), frontal parietal cingulum (C_FP), parahippocampal cingulum (C_PH), parahippocampal parietal cingulum (C_PHP), parolfactory cingulum (C_PO), corticospinal tract (CST), anterior corticostriatal tract (CS_A), posterior corticostriatal tract (CS_P), superior corticostriatal tract (CS_S), frontal aslant tract (FAT), inferior fronto-occipital fasciculus (IFOF), inferior longitudinal fasciculus (ILF), optic radiation (OR), parietal aslant tract (PAT), superior longitudinal fasciculus (SLF1), superior longitudinal fasciculus 2 (SLF2), superior longitudinal fasciculus 3 (SLF3), anterior thalamic radiation (TR_A), posterior thalamic radiation (TR_P), superior thalamic radiation (TR_S), uncinate fasciculus (UF), vertical occipital fasciculus (VOF), arcuate fasciculus (AF). See Supp. Table 7 for the full list of white matter tract name abbreviations.

Second, despite some inter-subject variability, the distribution of all subjects’ top-decoding subnetworks revealed population-level common encoding patterns. Specifically, we found that some regions were more frequently selected across subjects than others. We computed the frequency of each region being selected as part of the top-decoding subnetworks across all cross-validation folds among the population (Fig. 4b and 4e; Supp. Fig. 24) and found that this selection frequency was positively correlated with single-region decoding performance (Fig. 4c and 4f; Supp. Fig. 24), confirming the decoding stability of these regions. Therefore, we estimated each region’s population-level encoding strength of valence/arousal by linearly combining its selection frequency and single-region decoding performance (see Methods). We defined a population-level set of key-encoding regions as the regions whose encoding strength for valence or arousal ranked top 50% among all available regions.

Third, we explored in detail whether certain regions jointly encoded the valence and arousal or exhibited a stronger encoding preference towards one than the other. We divided the key-encoding regions into three groups according to each region’s relative encoding strength: (1) a shared group (comparable strong valence and arousal encoding), (2) a valence-preferred group (relatively stronger valence encoding), and (3) an arousal-preferred group (relatively stronger arousal encoding) (Fig. 4g). Our results showed that the shared group mainly included mesolimbic structures such as the amygdala (AMYG), hippocampus (HIP), insula (INS), lateral/medial orbitofrontal cortex (lOFC and mOFC), their densely connected temporal cortex (MTG and FFG), and fronto-parietal regions (rostral middle frontal cortex, RMF, and precuneus, PCUN). The valence-preferred group involved more posterior/lateral parts of the frontal and parietal cortices, including the inferior frontal gyrus (IFG), the caudal middle frontal gyrus (CMF), and the inferior parietal lobe (IPL), as well as the caudal anterior cingulate (cACC) cortex. The arousal-preferred group consisted of more posterior part of the cingulate (isthmus cingulate, IC), the primary somatosensory cortex (the postcentral gyrus, PoG), and especially, the thalamus (THA) (Fig. 4g and Supp. Fig. 26). Critically, these are typical regions that have been implicated in emotion processing by extensive neuroimaging and animal studies, showing the strong biological consistency of our encoding explanation (see Discussion).

The same analytical pipeline identified the white matter tract distribution of key-encoding signals (Fig. 4h-l; Supp. Fig. 25; Supp. Note 5), also revealing shared and preferred patterns for encoding (Fig. 4n; Supp. Fig. 27). The shared group mainly distributed over tracts that connect within limbic structures and connect limbic structures to widespread cortical regions. For example, the parahippocampal cingulum (C_PH), as the core white matter pathway of the limbic system, connects the anterior/posterior cingulate cortex and the parahippocampal gyrus; the inferior longitudinal fasciculus (ILF) and uncinate fasciculus (UF) connect the amygdala and hippocampus to widespread temporal, frontal, and occipital regions. Also, the arcuate fasciculus (AF) is the key tract bundle that directly connects the frontal and parietal lobes and forms the structural basis of the fronto-parietal network. The valence-preferred group mainly distributed over the parahippocampal parietal cingulum (C_PHP), connecting the limbic system to more localized posterior parts of the parietal cortex, and the anterior corticostriatal tract (CS_A) and superior longitudinal fasciculus II (SLF2), widely connecting the frontal regions to the striatum and parietal regions. The arousal-preferred group especially distributed over the anterior thalamic radiation (TR_A), connecting the thalamus to prefrontal and parietal regions, and the superior longitudinal fasciculus I (SLF1) and posterior corticostriatal tract (CS_P), connecting the postcentral gyrus to limbic, parietal and frontal regions.

### Spectral and temporal interpretation

We further used our neural decoding models to explain how spectral and temporal neural dynamics facilitate emotion decoding. In terms of single spectral band decoding, the performance using the *γ* band features consistently outperformed the other lower frequency bands (FDR-corrected *P* < 0.043; Supp. Figs. 28 and 29), further confirmed by a time-frequency analysis showing that the *γ* power significantly correlated with valence/arousal ratings across all trials and subjects (FDR-corrected cluster-based *P* < 0.042; Supp. Figs. 28 and 29). Moreover, the decoding performance when using all six frequency bands was significantly higher than using each single band alone (FDR-corrected *P* < 0.008; Supp. Figs. 28 and 29), suggesting complementary emotion-related information may be encoded by different frequency bands.

Our computational model allows for a detailed investigation of the temporal neural feature dynamics contributing to decoding. We quantified the timescale of decoded time-series for each subject by computing its frequency-domain power density (FPD, quantifying how fast the time-series varies over time, see Methods). Across the population, the FPD of decoded emotion time-series exceeded that of random-decoded time-series at timescales >16 s in both tasks (FDR-corrected *P <* 0.032 at each timescale; Supp. Fig. 30), indicating neural feature dynamics slower than tens of seconds mainly contributed to emotion decoding. The peaking FPD value roughly aligned with the period of showing the emotion-eliciting material in these two tasks, further confirming our identification of emotion-related temporal neural feature dynamics.

### Online implementation of neural decoding models and robust real-time emotion decoding

We finally implemented the neural decoding model online and demonstrated a robust real-time emotion decoding system in 4 new subjects (Fig. 5a; Supp. Tables 1,3,4,5 and 8). The online decoding system comprised: (1) an emotion-eliciting task presentation computer, (2) a camera for continuous and synchronized monitoring of the subject’s facial expression, (3) a neural interface device for real-time iEEG signal streaming, and (4) a laptop workstation dedicated to signal processing, neural decoding model implementation, and real-time decoding result visualization. For each subject, four personalized decoding models (video-task valence/arousal, image-task valence/arousal) were trained one day prior in a separate four-round experiment. Each model was then fixed and implemented in the online experiment (three rounds), respectively (Fig. 5b and 5c). Note that the subject was not informed of the neural signals being decoded or given feedback; the online system only passively decoded the emotion states and computed the online decoding performance measure. The measure, denoted as onlineCC, was defined as the CC between model-predicted valence/arousal ratings and the ground-truth scores rated up to the current time point. Note that the ground-truth scores were ensured to be unseen by the neural decoding model (see Methods).

**Fig. 5.**
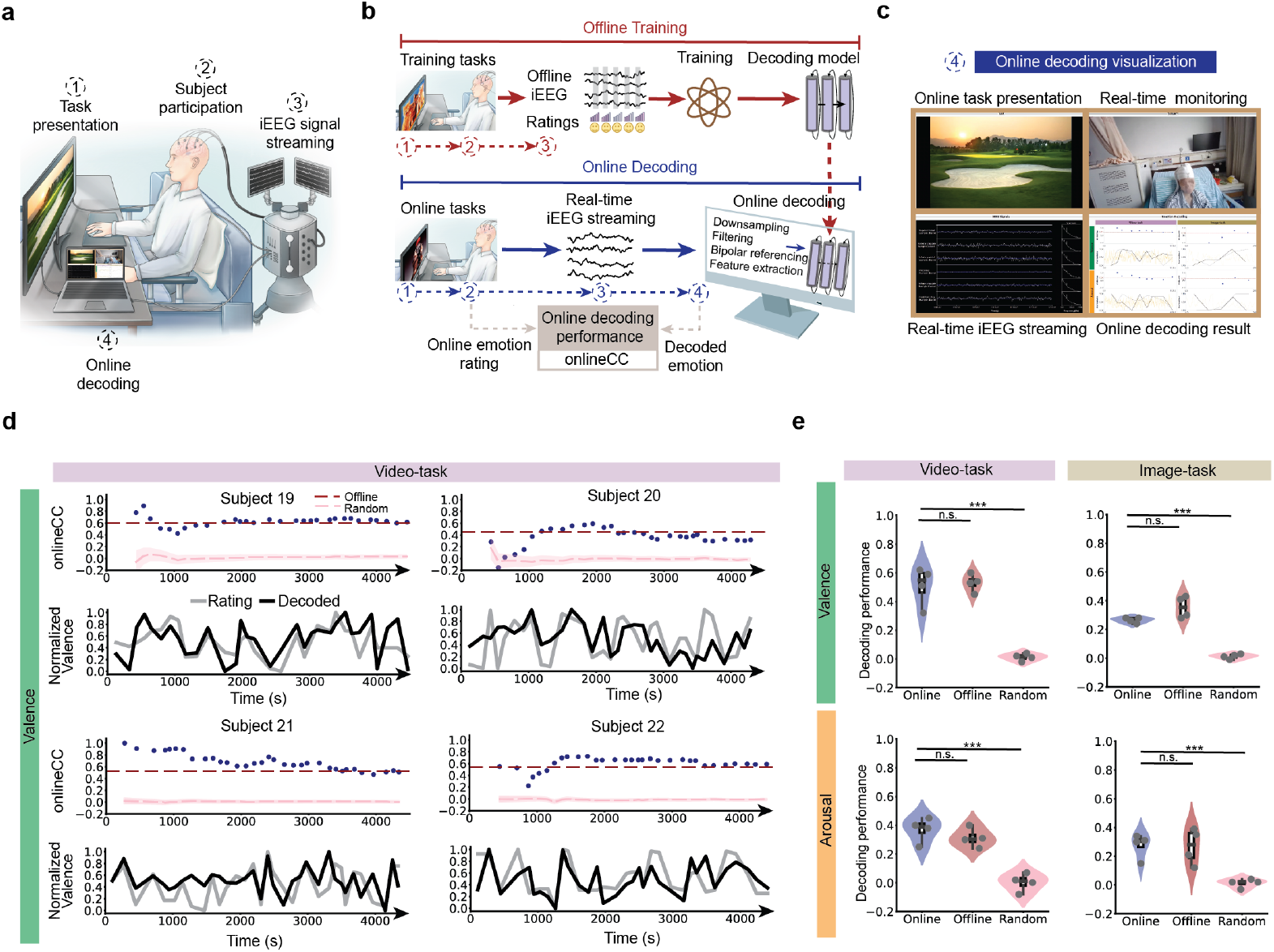
Online implementation of neural decoding models and robust real-time emotion decoding. **a**, The online emotion decoding experiment setup, which involves four key components as indicated by dotted circles. **b**, In each subject, the neural decoding model is trained in a separate offline training experiment (top) one day before the online decoding experiment (bottom). **c**, The online decoding visualization software streams the online task presentations, monitors the subject’s facial expressions, displays the real-time iEEG signals, and shows the online decoding results, which are all synchronized (see Supp. Video 1). **d**, The online valence decoding performance for each subject in the video-task (see Supp. Figs. 31-33 for other conditions). Top: the onlineCC values over time. The blue points indicate the onlineCC computed at each time point once the subject rated a valence score. Note that the onlineCC was updated by computing the correlation coefficient between all completed online ratings and decoding values up to the current time point. The onlineCC became stable after completing around 10 ratings, where enough samples had been accumulated for computing a reasonable correlation coefficient. The red dotted lines show the cvCC achieved by the offline training data. The pink dotted lines display the cvCC achieved by the offline training data with permuted labels, serving as the chance-level. Bottom: the decoded valence traces shown at each time point when the subject rated a valence score during the online experiments. **e**, The population pooled decoding performance comparison of online decoding, offline training, and chance-level. Each grey dot represents a subject (*N*=4).

In terms of the video-task, our real-time system enabled online valence and arousal decoding in each subject, with final onlineCC ranging from 0.32 to 0.62 and 0.25 to 0.45, respectively — significantly above chance level (*N* = 4, *n* = 32 to 34 per subject; FDR-corrected *P* < 0.05 for each subject; Fig. 5d; Supp. Fig. 31). The real-time decoded traces of valence and arousal closely followed the ground-truth ratings (Fig. 5d). The population-pooled onlineCC reached 0.51±0.07 and 0.37±0.04 for valence and arousal, respectively (*n* = 132, *P* < 0.001; Fig. 5e), comparable to offline training performance (*n* = 182, cvCC = 0.53±0.03 and 0.31±0.03 for valence and arousal, respectively; t-test *P* > 0.06 in both cases). Similar results held for online image-task decoding (Fig. 5e; Supp. Figs. 32 and 33). The average latency was 376±11.87 milliseconds across all tasks, subjects, and trials (*n* = 564, see Supp. Table 11 for computational complexity and time). These results demonstrate low-latency, high-performance online decoding of emotion states from iEEG.

## Discussion

We developed personalized neural decoding models that effectively decoded fine-grained continuous values of the two major dimension of emotion (valence and arousal), whose performance was boosted by integrating grey and white matter iEEG network signals. The model more than doubled the R-squared performance of prior EEG and iEEG emotion decoding studies. Further, our decoding was interchangeable and transferable across two distinct emotion-eliciting tasks, suggesting common neural representations of emotion states elicited by different stimuli. Moreover, the decoding models further revealed shared and preferred mesolimbic-thalamo-cortical subnetworks encoding valence and arousal, showing neurophysiological explainability. Finally, we implemented the models online and demonstrated robust real-time emotion decoding. Together, these advancements form an integrated computational framework for intracranial neural activity-based human emotion decoding, explanation, and application.

We used task-based emotion-eliciting paradigms to elicit standardized, reproducible, rapid and controlled emotion responses that allowed for the collection of a much larger number of emotion state measurements than spontaneous emotion assessment paradigms^79,80^. In terms of stimulus type selection, compared to alternative methods (e.g., recall^81^ and hypnosis^82^), audiovisual stimuli offer better standardization and reproducibility^80^. The video-task, with audiovisual inputs and narrative engagement, evokes stronger time-varying responses^83^, whereas the image-task allows more standardized and faster emotion elicitation^72^. Overall, the quantity, quality, and diversity of our data enabled high-performance cross-task decoding.

In terms of decoding performance, our model surpassed several related lines of studies. First, several recent iEEG investigations into the spectral dynamics of emotion processing have reported modest linear correlation coefficients between spectral features and valence/arousal scores^35,36^ (on average 0.42±0.02, Supp. Table 12) from an encoding perspective, pointing to both the possibility and challenge of decoding nuanced dynamics of emotion. Second, a series of recent studies have used iEEG to directly decode mood states or depression symptoms with rigorous cross-validation^40–46,59^, with a cohort-size-weighted average cvCC=0.57±0.02 (Supp. Table 13). Comparing our decoding performance (e.g., video-task valence cvCC of 0.70±0.02) to these studies should take caution due to the differences in experimental design, behavioral measurements, and temporal resolution. Task-evoked emotions may exhibit larger and more obvious variations than naturalistic mood fluctuations; on the other hand, our emotion measurement is a single-item rating of valence or arousal, potentially introducing more noise compared to the multi-item questionnaires typically employed in mood or symptom assessments (e.g., Immediate Mood Scaler^84^ and Computerized Adaptive Test–Depression Inventory^85^). Mood decoding captures more stable, enduring affective states; by contrast, we decoded more rapid emotion fluctuations, providing distinct insights into the faster dynamics of affect processing as revealed by our temporal interpretation analyses. Third, in terms of the coarse-grained discrete emotion classification, our three-class cvACC of 0.75 (two-class cvACC of 0.81) outperformed previous iEEG emotion classification studies with three-class cvACC being up to 0.68^39^ (two-class cvACC of 0.73^38^ and 0.62^37^). Finally, consistent with prior studies^11,17^, *γ* band mainly contributed to our decoding, suggesting high-performance emotion decoding may require intensive use of intracranial signals given the limited capacity of EEG and fMRI in capturing high-frequency neural dynamics. Overall, the above comparisons show the high performance of our neural decoding models.

Regarding cross-task decoding, previous iEEG and EEG studies focused on cross-time/cross-subject generalization, all within a single task^11–13,17,21–23^, which did not investigate or sufficiently leverage the central integrative nature of emotion states across diverse tasks. Our capability of interchangeably decoding suggests common neural representations of emotion states. Our transferable decoding not only reduced the risk of overfitting by leveraging more data but also indicated the practical utility for ecological emotion applications requiring rapid deployment across diverse contexts^67–69^.

Real-time emotion decoding provides a translational foundation for next-generation engineering and clinical applications, but prior studies are largely limited to offline, with only a few recent DBS studies achieving real-time computation of single-channel, single-band power feature that correlates with affective disorder symptoms^34,59,86^. Our work moved a step forward by realizing online decoding, which has the potential of providing real-time feedback for guiding interventions of disorders. Also, our task-based paradigm naturally fits into the neurofeedback training framework^87^. Our focus on intracranial decoding closely relates to the aDBS therapy for affective disorders such as treatment-resistant depression (TRD)^59^ and post-traumatic stress disorder (PTSD)^34^. aDBS aims to use invasive local field potentials to decode naturalistic or event-triggered affective symptom variations as real-time feedback to guide stimulation delivery^67–69,88^. Our transferable framework can provide useful ‘pre-trained’ models further optimized for such applications. Moreover, chronic applications such as aDBS require the temporal stability of decoding over days to weeks^67–69^; under the practical constraint of our data collection window, we showed that emotion decoding remained temporally stable for up to 44 hours, providing initial evidence for its practical use. Note that the offline training performance from our online experiments was lower than our main offline experiments because the online recording device had fewer channels and larger noise (Supp. Fig. 34). Thus, incorporating our systems with neurofeedback training and aDBS are promising directions but require further optimization of accuracy, stability, and latency (see Supp. Note 6).

Our entire offline and online decoding framework uses the primary analytical approach of training and validating a personalized valence decoding model and a personalized arousal decoding model for each single subject, which forms the basis of high-performance within-subject decoding in practical applications. Fitting a common population-level decoding model by aggregating data from all subjects faces the critical challenge of electrode implantation and data distribution heterogeneity, which may be addressed by large language model (LLM)-inspired brain signal foundation models^89^ in future research. Nevertheless, from a neural encoding explainability perspective, we proposed a secondary population-level analytical approach of aggregating decoding results across the individually-fitted personalized models to statistically investigate common encoding patterns across subjects. Such population-level analyses can advance the understanding of the general emotion encoding mechanisms and serve as a priori guidance on electrode placement in new subjects, as we expand the discussion below.

Recent intracranial research has advanced understanding of the distributed mood/emotion networks^11,12,17^, yet remains limited by cohort size, coverage, and decoding accuracy— impeding fine-grained subnetwork and dimensional specificity analysis. Our study partially closed the gap by using our decoding models to reveal that subnetworks of roughly four regions sufficed for decoding, consistent with findings that small subnetworks can encode emotion, mood, and symptoms^10,11,17,90^ and carefully-designed computational methods can efficiently aggregate such information for decoding^39,43–46,59^. The spatial distribution of these key subnetworks varied across subjects, likely due to implantation differences and possible heterogeneity in affect-regulating circuits—consistent with recent findings of varied key regions even under consistent implantations^43^.

Nevertheless, our population-level analytical approach revealed that common mesolimbic-thalamo-cortical regions differentially encoded valence and arousal. The shared group mainly consisted of mesolimbic structures (AMYG, HIP, INS, lOFC, mOFC)—a well-known core emotion processing network^91,92^, and the fronto-parietal regions (RMF, PCUN) for task-directed, domain-general cognitive control of emotion regulation^93–95^; alternations of these core circuits have been reported across various affective disorders such as depression and anxiety^90^. The valence-preferred group included more posterior/lateral frontal-parietal areas (CMF, IFG, IPL)— related to social valence^96^, and the anterior cingulate cortex (cACC)— processing valence-specific information about value and uncertainty^97^. The arousal-preferred group highlighted the thalamus—critical in awake-sleep^98^ and consciousness regularization^99^, the isthmus cingulate cortex (IC)— key structure for arousal and awareness^100^, and the postcentral gyrus, known for somatosensory information processing^101^. These findings together provide neurophysiological explainability for our decoding models and delineate a detailed spatial distribution of key regions supporting valence and arousal processing.

Our results highlight the importance of white matter in emotion decoding by expanding conventional grey matter-focused approaches. Conventional approaches^35–59^ have typically treated white matter signals as noise to reduce neural feature complexity. However, in theory, white matter signals can contain useful information as they represent a mixture of activities from nearby and distant grey matter^102^. Consequently, while increasingly studied in fMRI^103,104^, white matter remains underexplored in iEEG, with only a few studies for decoding movements or speech^105–108^. Given our dataset scale and model design, we demonstrated that white matter signals effectively decoded emotion and boosted performance when combined with grey matter. However, it should be noted that the exact source of white matter iEEG signals is still under investigation. A recent study has suggested that white matter iEEG signals can reflect the information transmission along white matter tracts themselves^109^. On the other hand, white matter signals can also be substantially influenced by volume conduction from distant sources^102^. Nevertheless, the white matter tract distribution of the contributing iEEG channels was concretely summarized by our population-level analytical approach. The key white matter tract distribution is linked to affective disorders in prior neuroimaging study^110^, mainly connecting the limbic system, thalamus, postcentral gyrus, and the fronto-parietal network—consistent with our identified grey matter regions and supporting the emerging theory of emotion as a distributed network process^111^. Whether such distribution directly characterizes the genuine neural activity along white matter tracts requires future research though.

Valence and arousal have long been considered as two major emotion primitives, but whether they are independent dimensions remains unknown. Although emotion theories often conceptualize them as distinct dimensions, there has been no direct evidence supporting that they are completely independent^112^. Indeed, previous empirical studies^72^ have consistently shown that self-rated valence and arousal scores exhibit a nonlinearly correlated “boomerang” pattern. In the context of emotion decoding, prior studies^35–59^ have rarely explored the relationship between decoded valence and arousal. Our decoding analyses reveal that even though we trained separate models for decoding valence and arousal, the decoded signals’ correlation pattern closely mirrored the self-rated “boomerang” distribution, demonstrating the behavioral consistency of our decoding framework and indicating that the two decoded signals provide both redundant and complementary information in a nonlinear manner. Consistently, our spatial analyses identified both shared and preferred subnetworks encoding valence and arousal. However, dissecting the neural mechanisms underlying valence-arousal interaction, quantifying the exact amount of their redundant and complementary information, and leveraging such information to further improve decoding all warrant further investigation.

The personalized decoding performance and population-level encoding explainability collectively provide some preliminary guidance on the conditions under which our decoding framework can be productively applied in practical scenarios. First, important implantation regions for the joint decoding of valence and arousal mainly spread mesolimbic and fronto-parietal areas, and for the specific decoding of valence and arousal mainly involve the inferior frontal gyrus and thalamus, respectively. Second, for a given subject, using signals spreading four regions may be enough for achieving a sufficiently high decoding performance, but the exact distribution of these four regions can be subject-dependent. Third, further down to the electrode and channel level, compared with the best-performing case of using all channels, the more computationally efficient cases of using best single-electrode and best single-channel resulted in a drop of cvCC by 0.11 and 0.10 for our model, respectively; the worse but still significant decoding of using simpler gradient boosting models asks for careful and tailored considerations of the trade-off between performance and complexity for specialized applications. Fourth, transfer decoding can be achieved across tasks that share similar valence-arousal evoking capability (e.g., the “boomerang” pattern), where a source-task trained decoding model can reach near-optimal target-task performance after fine-tuning with on average 43 trials in our case (50% of target-task training trials). Finally, online deployment requires a low-latency streaming and processing pipeline (around 376ms latency in our case), where the total delay must be less than the decoding step size (one second in our case) for supporting real-time applications; such a pipeline demands hardware capability of fast data access, optimized signal synchronization, along with efficient neural feature extraction and model inference.

Our work has several limitations. First, our cohort consisted exclusively of subjects with epilepsy; while our control analyses excluded the influence of epileptic activity on decoding, the subjects’ pathological brain network and electrode implantation heterogeneity can still influence the interpretability of the results. Future studies could recruit patients with primary affective disorders and use consistent electrode implantations^37,40,43,113,114^. Second, we only included two emotion-eliciting tasks; validation across a broader range of emotion-eliciting methods, e.g., imagined emotions^81^, would enhance generalizability. Third, we decoded and interpreted valence and arousal as the major dimension of emotion, but other dimension (e.g., dominance and expectation^115,116^) should be considered in the future to formulate a multi-dimensional computational framework. Fourth, we only considered the widely-used spectral neural features as model input, but time-domain features may provide substantial complementary information to improve decoding, especially when channel numbers are limited^37,113^; investigating how frequency-domain and time-domain features jointly encode emotion and synergistically decode emotion is a critical future direction. Finally, while our subject cohort size and electrode coverage were relatively large compared to previous iEEG mood and emotion decoding studies, future work should consider incorporating more subjects (especially online), augmenting coverage by simultaneously recording scalp EEG, and improving signal resolution by simultaneously recording spiking activity^117,118^ to further improve performance and explainability.

## Outlook

Taken as a whole, our results demonstrate high-performance, task-transferable, explainable, and real-time decoding of human emotion states from intracranial neural activity, which can advance emotion decoding neurotechnology toward deployable affective brain–computer interfaces and closed-loop therapeutic systems for affective disorders.

## Data availability

The main data supporting the results in this study are available within the paper and its Supplementary Information. The collected raw neural and behavioural data are a modified version of clinical recordings for the purpose of seizure localization and clinical decisions. Thus, the minimum de-identified dataset used to generate the findings of this study will be available upon reasonable request to the corresponding author. Processed data for generating the figures will be available upon publication.

## Code availability

The iEEG data preprocessing was implemented by MATLAB 2020b custom scripts mainly relying on the FieldTrip toolbox. The decoding model construction, fitting, and evaluation were implemented by Python 3.0 custom scripts. All codes will be available upon publication.

## Acknowledgements

This work was supported in part by the National Natural Science Foundation of China under Grants 62306269, 62476239, and 62336007, the National Key Research and Development Program of China under Grant 2023YFC2506200, the Key R&D Program of Zhejiang under Grant 2025C01112, the Zhejiang Provincial Natural Science Foundation of China under Grant LD24H090001, and the Nanhu Brain-computer Interface Institute under Grant 010901008.

## Author contributions

Y.Y., S.H., J.Z. and Y.W. supervised the project. Y.Y. and Y.W. conceived the emotion decoding framework, and Y.Y. and W.C. developed it. Y.Y., W.C., and S.H. designed the experiment. W.C. and Y.C. implemented the experimental system. W.C., Y.C., L.D., and C.Z. performed the experiments and data collection. H.J., Z.Z., X.G., and S.W. identified the electrode position using imaging data and assessed the neurological conditions of the subjects. N.W. and S.H. performed the cognitive and psychiatric assessment. Y.Y. and W.C. implemented and performed the modeling and analyses. Y.Y. and Y.W. supervised all the modelling and analysis work. S.H. and J.Z. supervised all clinical aspects of the study. Y.Y. and W.C. wrote the manuscript with input from Y.W., S.H., J.Z., and G.P.

## References

1. Bach, D. R. & Dayan, P. Algorithms for survival: a comparative perspective on emotions. Nat. Rev. Neurosci. 18, 311–319 (2017).

2. Lincoln, T. M., Schulze, L. & Renneberg, B. The role of emotion regulation in the characterization, development and treatment of psychopathology. Nat. Rev. Psychol. 1, 272–286 (2022).

3. Ekman, P. An argument for basic emotions. Cogn. Emot. 6, 169–200 (1992).

4. Panksepp, J. Affective Neuroscience: The Foundations of Human and Animal Emotions. (Oxford university press, 2004).

5. Schachter, S. & Singer, J. E. Cognitive, social, and physiological determinants of emotional state. Psychol. Rev. 69, 379–399 (1962).

6. Arnold, M. B. Emotion and Personality. (1960).

7. Russell, J. A. A circumplex model of affect. J. Pers. Soc. Psychol. 39, 1161–1178 (1980).

8. Barrett, L. F. The theory of constructed emotion: an active inference account of interoception and categorization. Soc. Cogn. Affect. Neurosci. 12, 1–23 (2016).

9. Anderson, D. J. & Adolphs, R. A Framework for Studying Emotions across Species. Cell 157, 187–200 (2014).

10. Malezieux, M., Klein, A. S. & Gogolla, N. Neural Circuits for Emotion. Annu. Rev. Neurosci. 46, 211–231 (2023).

11. Kabotyanski, K. E., Provenza, N. R. & Sheth, S. A. Intracranial neural biomarkers of psychiatric symptoms and their utility for guiding neuromodulation therapy: a systematic review. Biol. Psychiatry https://doi.org/10.1016/j.biopsych.2025.10.005 (2025) doi:10.1016/j.biopsych.2025.10.005.

12. Shanechi, M. M. Brain–machine interfaces from motor to mood. Nat Neurosci 22, 1554–1564 (2019).

13. Lim, R. Y., Lew, W.-C. L. & Ang, K. K. Review of EEG Affective Recognition with a Neuroscience Perspective. Brain Sci. 14, 364 (2024).

14. Kragel, P. A. & LaBar, K. S. Decoding the Nature of Emotion in the Brain. Trends Cogn. Sci. 20, 444–455 (2016).

15. Rossi, S. Emotion Recognition for Human-Robot Interaction: Recent Advances and Future Perspectives. Front. Robot. AI 7, (2020).

16. Scangos, K. W., State, M. W., Miller, A. H., Baker, J. T. & Williams, L. M. New and emerging approaches to treat psychiatric disorders. Nat. Med. 29, 317–333 (2023).

17. Ramos Lopez, C. G. et al. Stereotactic electroencephalography in epilepsy patients for mapping of neural circuits related to emotional and psychiatric behaviors: a systematic review. Neurosurg. Focus 54, E4 (2023).

18. Siedlecka, E. & Denson, T. F. Experimental Methods for Inducing Basic Emotions: A Qualitative Review. Emot. Rev. 11, 87–97 (2019).

19. Underwood, R., Tolmeijer, E., Wibroe, J., Peters, E. & Mason, L. Networks underpinning emotion: A systematic review and synthesis of functional and effective connectivity. NeuroImage 243, 118486 (2021).

20. Logothetis, N. K. What we can do and what we cannot do with fMRI. Nature 453, 869–878 (2008).

21. Alarcao, S. M. & Fonseca, M. J. Emotions Recognition Using EEG Signals: A Survey. IEEE Trans. Affect. Comput. 10, 374–393 (2019).

22. Wu, D., Lu, B.-L., Hu, B. & Zeng, Z. Affective Brain–Computer Interfaces (aBCIs): A Tutorial. Proc. IEEE 111, 1314–1332 (2023).

23. Wu, D., Xu, Y. & Lu, B.-L. Transfer Learning for EEG-Based Brain–Computer Interfaces: A Review of Progress Made Since 2016. IEEE Trans. Cogn. Dev. Syst. 14, 4–19 (2022).

24. Zhang, S., Tang, C. & Guan, C. Visual-to-EEG cross-modal knowledge distillation for continuous emotion recognition. Pattern Recognit. 130, 108833 (2022).

25. Ding, Y. & Guan, C. GIGN: Learning Graph-in-graph Representations of EEG Signals for Continuous Emotion Recognition.

26. Ding, Y., Zhang, S., Tang, C. & Guan, C. MASA-TCN: Multi-Anchor Space-Aware Temporal Convolutional Neural Networks for Continuous and Discrete EEG Emotion Recognition. IEEE J. Biomed. Health Inform. 28, 3953–3964 (2024).

27. Boucher, O. et al. Spatiotemporal dynamics of affective picture processing revealed by intracranial high-gamma modulations: Intracranial EEG and Affective Pictures. Hum. Brain Mapp. 36, 16–28 (2015).

28. Qasim, S. E., Mohan, U. R., Stein, J. M. & Jacobs, J. Neuronal activity in the human amygdala and hippocampus enhances emotional memory encoding. Nat. Hum. Behav. 7, 754–764 (2023).

29. Kauvar, I. et al. Conserved brain-wide emergence of emotional response from sensory experience in humans and mice. Science 388, eadt3971 (2025).

30. Fan, X. et al. Brain mechanisms underlying the emotion processing bias in treatment-resistant depression. Nat. Ment. Health 2, 583–592 (2024).

31. Pan, N. C. et al. Left Insula and Right Middle Temporal Gyrus Dominate Cortical Network Discriminating Arousal-dependent Emotions. Adv. Sci. 12, 2411790 (2025).

32. Metzger, B. A. et al. Intracranial stimulation and EEG feature analysis reveal affective salience network specialization. Brain 146, 4366–4377 (2023).

33. Buot, A. et al. Emotions Modulate Subthalamic Nucleus Activity: New Evidence in Obsessive-Compulsive Disorder and Parkinson’s Disease Patients. Biol. Psychiatry Cogn. Neurosci. Neuroimaging 6, 556–567 (2021).

34. Gill, J. L. et al. A pilot study of closed-loop neuromodulation for treatment-resistant post-traumatic stress disorder. Nat. Commun. 14, 2997 (2023).

35. Huebl, J. et al. Processing of emotional stimuli is reflected by modulations of beta band activity in the subgenual anterior cingulate cortex in patients with treatment resistant depression. Soc. Cogn. Affect. Neurosci. 11, 1290–1298 (2016).

36. Sonkusare, S. et al. Frequency dependent emotion differentiation and directional coupling in amygdala, orbitofrontal and medial prefrontal cortex network with intracranial recordings. Mol. Psychiatry 28, 1636–1646 (2023).

37. Merk, T. et al. Invasive neurophysiology and whole brain connectomics for neural decoding in patients with brain implants. Nat. Biomed. Eng. 1–18 (2025) doi:10.1038/s41551-025-01467-9.

38. Azadian, E. et al. Decoding Happiness from Neural and Video Recordings for Better Insight Into Emotional Processing in the Brain. in 2021 43rd Annual International Conference of the IEEE Engineering in Medicine & Biology Society (EMBC) 6747–6750 (IEEE, Mexico, 2021). doi:10.1109/EMBC46164.2021.9629972.

39. Bijanzadeh, M. et al. Decoding naturalistic affective behaviour from spectro-spatial features in multiday human iEEG. Nat. Hum. Behav. 6, 823–836 (2022).

40. Alagapan, S. et al. Cingulate dynamics track depression recovery with deep brain stimulation. Nature 622, 130–138 (2023).

41. Frank, A. et al. Identification of a personalized intracranial biomarker of depression and response to DBS therapy. Brain Stimulat. 16, 383 (2021).

42. Cao, D. State-specific modulation of mood using intracranial electrical stimulation of the orbitofrontal cortex. Brain Stimulat. 16, 1112–1122 (2023).

43. Xiao, J. et al. Decoding Depression Severity From Intracranial Neural Activity. Biol. Psychiatry 94, 445–453 (2023).

44. Sani, O. G. et al. Mood variations decoded from multi-site intracranial human brain activity. Nat Biotechnol 36, 954–961 (2018).

45. Kirkby, L. A. et al. An Amygdala-Hippocampus Subnetwork that Encodes Variation in Human Mood. Cell 175, 1688–1700.e14 (2018).

46. Scangos, K. W. et al. Pilot Study of An Intracranial Electroencephalography Biomarker of Depressive Symptoms in Epilepsy. J. Neuropsychiatry Clin. Neurosci. 32, 185–190 (2020).

47. Voon, V. et al. Bed Nucleus of the Stria Terminalis-Nucleus Accumbens Deep Brain Stimulation for Depression: A Randomized Controlled Trial and an Intracranial Physiological Biomarker Predictor. Preprint at 10.21203/rs.3.rs-4854344/v1 (2024).

48. Veerakumar, A. et al. Field potential 1/*f* activity in the subcallosal cingulate region as a candidate signal for monitoring deep brain stimulation for treatment-resistant depression. J. Neurophysiol. 122, 1023–1035 (2019).

49. Sendi, M. S. E. et al. Intraoperative neural signals predict rapid antidepressant effects of deep brain stimulation. Transl. Psychiatry 11, 551 (2021).

50. Iravani, B. et al. Intracranial Recordings of the Human Orbitofrontal Cortical Activity during Self-Referential Episodic and Valenced Self-Judgments. J. Neurosci. 44, e1634232024 (2024).

51. Hacker, C. et al. Aperiodic (1/f) Neural Activity Robustly Tracks Symptom Severity Changes in Treatment-Resistant Depression. Biol. Psychiatry Cogn. Neurosci. Neuroimaging 10, 186–194 (2025).

52. Sonkusare, S. et al. Power signatures of habenular neuronal signals in patients with bipolar or unipolar depressive disorders correlate with their disease severity. Transl. Psychiatry 12, 72 (2022).

53. Clark, D. L., Brown, E. C., Ramasubbu, R. & Kiss, Z. H. T. Intrinsic Local Beta Oscillations in the Subgenual Cingulate Relate to Depressive Symptoms in Treatment-Resistant Depression. Biol. Psychiatry 80, e93–e94 (2016).

54. Neumann, W.-J. et al. Different patterns of local field potentials from limbic DBS targets in patients with major depressive and obsessive compulsive disorder. Mol. Psychiatry 19, 1186–1192 (2014).

55. Merkl, A. et al. Modulation of Beta-Band Activity in the Subgenual Anterior Cingulate Cortex during Emotional Empathy in Treatment-Resistant Depression. Cereb. Cortex 26, 2626–2638 (2016).

56. De Hemptinne, C. et al. Prefrontal Physiomarkers of Anxiety and Depression in Parkinson’s Disease. Front. Neurosci. 15, 748165 (2021).

57. Rao, V. R. et al. Direct Electrical Stimulation of Lateral Orbitofrontal Cortex Acutely Improves Mood in Individuals with Symptoms of Depression. Curr. Biol. 28, 3893–3902.e4 (2018).

58. Zhang, C. et al. Bilateral Habenula deep brain stimulation for treatment-resistant depression: clinical findings and electrophysiological features. Transl. Psychiatry 12, 52 (2022).

59. Scangos, K. W. et al. Closed-loop neuromodulation in an individual with treatment-resistant depression. Nat. Med. 27, 1696–1700 (2021).

60. Qasim, S. E., Deswal, A., Saez, I. & Gu, X. Positive affect modulates memory by regulating the influence of reward prediction errors. Commun. Psychol. 2, 52 (2024).

61. Landré, E., Chipaux, M., Maillard, L., Szurhaj, W. & Trébuchon, A. Electrophysiological technical procedures. Neurophysiol. Clin. 48, 47–52 (2018).

62. Zhang, S. et al. Disruption of superficial white matter in the emotion regulation network in bipolar disorder. NeuroImage Clin. 20, 875–882 (2018).

63. Forkel, S. J., Friedrich, P., Thiebaut De Schotten, M. & Howells, H. White matter variability, cognition, and disorders: a systematic review. Brain Struct. Funct. 227, 529–544 (2022).

64. Silversmith, D. B. et al. Plug-and-play control of a brain–computer interface through neural map stabilization. Nat. Biotechnol. 39, 326–335 (2021).

65. Spitale, M. et al. Past, Present, and Future: A Survey of The Evolution of Affective Robotics For Well-being. IEEE Trans. Affect. Comput. 1–17 (2025).

66. Fernández-Alvarez, J. et al. Efficacy of bio- and neurofeedback for depression: a meta-analysis. Psychol. Med. 52, 201–216 (2022).

67. Oganesian, L. L. & Shanechi, M. M. Brain–computer interfaces for neuropsychiatric disorders. Nat. Rev. Bioeng. 2, 653–670 (2024).

68. Sellers, K. K. et al. Closed-loop neurostimulation for the treatment of psychiatric disorders. Neuropsychopharmacology 49, 163–178 (2024).

69. Herron, J. et al. Challenges and opportunities of acquiring cortical recordings for chronic adaptive deep brain stimulation. Nat. Biomed. Eng. 9, 606–617 (2024).

70. Johnson, K. A., Okun, M. S., Scangos, K. W., Mayberg, H. S. & De Hemptinne, C. Deep brain stimulation for refractory major depressive disorder: a comprehensive review. Mol. Psychiatry 29, 1075–1087 (2024).

71. Oehrn, C. R. et al. Chronic adaptive deep brain stimulation versus conventional stimulation in Parkinson’s disease: a blinded randomized feasibility trial. Nat. Med. 30, 3345–3356 (2024).

72. Branco, D., Gonçalves, Ó. F. & Badia, S. B. I. A Systematic Review of International Affective Picture System (IAPS) around the World. Sensors 23, 3866 (2023).

73. Klein, A. & Tourville, J. 101 Labeled Brain Images and a Consistent Human Cortical Labeling Protocol. Front. Neurosci. 6, 33392 (2012).

74. Fischl, B. et al. Whole Brain Segmentation: Automated Labeling of Neuroanatomical Structures in the Human Brain. Neuron 33, 341–355 (2002).

75. Van Essen, D. C. et al. The Human Connectome Project: A data acquisition perspective. NeuroImage 62, 2222–2231 (2012).

76. Schneider, S., Lee, J. H. & Mathis, M. W. Learnable latent embeddings for joint behavioural and neural analysis. Nature 617, 360–368 (2023).

77. Gunnarsdottir, K. M. Source-sink connectivity: a novel interictal EEG marker for seizure localization. Brain 145, 3901–3915 (2022).

78. Bentéjac, C., Csörgő, A. & Martínez-Muñoz, G. A comparative analysis of gradient boosting algorithms. Artif. Intell. Rev. 54, 1937–1967 (2021).

79. Ciuk, D., Troy, A. K. & Jones, M. C. Measuring Emotion: Self-Reports vs. Physiological Indicators. SSRN Electron. J. (2015).

80. Gerdes, A. B. M., Wieser, M. J. & Alpers, G. W. Emotional pictures and sounds: a review of multimodal interactions of emotion cues in multiple domains. Front. Psychol. 5, 1351 (2014).

81. Chanel, G., Kierkels, J. J. M., Soleymani, M. & Pun, T. Short-term emotion assessment in a recall paradigm. Int. J. Hum.-Comput. Stud. 67, 607–627 (2009).

82. Houghton, L. A. Visceral sensation and emotion: a study using hypnosis. Gut 51, 701–704 (2002).

83. Uhrig, M. K. et al. Emotion Elicitation: A Comparison of Pictures and Films. Front. Psychol. 7, 180 (2016).

84. Nahum, M. et al. Immediate Mood Scaler: Tracking Symptoms of Depression and Anxiety Using a Novel Mobile Mood Scale. JMIR MHealth UHealth 5, e6544 (2017).

85. Gibbons, R. D. et al. Development of a Computerized Adaptive Test for Depression. Arch. Gen. Psychiatry 69, 1104 (2012).

86. Nho, Y.-H. et al. Responsive deep brain stimulation guided by ventral striatal electrophysiology of obsession durably ameliorates compulsion. Neuron 112, 73–83.e4 (2024).

87. Sitaram, R. et al. Closed-loop brain training: the science of neurofeedback. Nat. Rev. Neurosci. 18, 86–100 (2017).

88. Yang, Y. et al. Modelling and prediction of the dynamic responses of large-scale brain networks during direct electrical stimulation. Nat Biomed Eng 5, 324–345 (2021).

89. Zhou, X. et al. Brain Foundation Models: A survey on advancements in neural signal processing and brain discovery. IEEE Signal Process. Mag. 42, 22–35 (2025).

90. Williams, L. M. Precision psychiatry: a neural circuit taxonomy for depression and anxiety. Lancet Psychiatry 3, 472–480 (2016).

91. Rolls, E. T. Limbic systems for emotion and for memory, but no single limbic system. Cortex 62, 119–157 (2015).

92. Manssuer, L. et al. Reward recalibrates rule representations in human amygdala and hippocampus intracranial recordings. Nat. Commun. 15, 9518 (2024).

93. Cole, M. W., Repovš, G. & Anticevic, A. The Frontoparietal Control System: A Central Role in Mental Health. The Neuroscientist 20, 652–664 (2014).

94. Taylor, C. et al. An Exploratory Study of Large-Scale Brain Networks during Gambling Using SEEG. Brain Sci. 14, 773 (2024).

95. Hoy, C. W. et al. Beta and theta oscillations track effort and previous reward in the human basal ganglia and prefrontal cortex during decision making. Proc. Natl. Acad. Sci. 121, e2322869121 (2024).

96. Padilla-Coreano, N., Tye, K. M. & Zelikowsky, M. Dynamic influences on the neural encoding of social valence. Nat. Rev. Neurosci. 23, 535–550 (2022).

97. Monosov, I. E. Anterior cingulate is a source of valence-specific information about value and uncertainty. Nat. Commun. 8, 134 (2017).

98. Gent, T. C., Bandarabadi, M., Herrera, C. G. & Adamantidis, A. R. Thalamic dual control of sleep and wakefulness. Nat. Neurosci. 21, 974–984 (2018).

99. Fang, Z. et al. Human high-order thalamic nuclei gate conscious perception through the thalamofrontal loop. Science 388, eadr3675 (2025).

100. Leech, R. & Sharp, D. J. The role of the posterior cingulate cortex in cognition and disease. Brain 137, 12–32 (2014).

101. Anders, S., Lotze, M., Erb, M., Grodd, W. & Birbaumer, N. Brain activity underlying emotional valence and arousal: A response-related fMRI study. Hum. Brain Mapp. 23, 200–209 (2004).

102. Mercier, M. R. et al. Evaluation of cortical local field potential diffusion in stereotactic electro-encephalography recordings: A glimpse on white matter signal. NeuroImage 147, 219–232 (2017).

103. Berluti, K., Ploe, M. L. & Marsh, A. A. Emotion processing in youths with conduct problems: an fMRI meta-analysis. Transl. Psychiatry 13, 105 (2023).

104. Seyedmirzaei, H. et al. White matter tracts associated with alexithymia and emotion regulation: A diffusion MRI study. J. Affect. Disord. 314, 271–280 (2022).

105. Li, G. et al. Detection of human white matter activation and evaluation of its function in movement decoding using stereo-electroencephalography (SEEG). J. Neural Eng. 18, 0460c6 (2021).

106. Bouton, C. et al. Decoding Neural Activity in Sulcal and White Matter Areas of the Brain to Accurately Predict Individual Finger Movement and Tactile Stimuli of the Human Hand. Front. Neurosci. 15, 699631 (2021).

107. Ottenhoff, M. C. et al. Decoding continuous goal-directed movement from human brain-wide intracranial recordings. Preprint at 10.1101/2025.02.05.636287 (2025).

108. Soroush, P. Z. et al. Contributions of Stereotactic EEG Electrodes in Grey and White Matter to Speech Activity Detection. in 2022 44th Annual International Conference of the IEEE Engineering in Medicine & Biology Society (EMBC) 4789–4792 (IEEE, Glasgow, Scotland, United Kingdom, 2022).

109. Revell, A. Y. et al. White Matter Signals Reflect Information Transmission Between Brain Regions During Seizures. Brain 149, 77–89 (2026).

110. Koshiyama, D. et al. White matter microstructural alterations across four major psychiatric disorders: mega-analysis study in 2937 individuals. Mol. Psychiatry 25, 883–895 (2020).

111. LeDuke, D. O., Borio, M., Miranda, R. & Tye, K. M. Anxiety and depression: A top-down, bottom-up model of circuit function. Ann. N. Y. Acad. Sci. 1525, 70–87 (2023).

112. Kuppens, P., Tuerlinckx, F., Russell, J. A. & Barrett, L. F. The relation between valence and arousal in subjective experience. Psychol. Bull. 139, 917–940 (2013).

113. Liu, S. et al. A Habenula Neural Biomarker Simultaneously Tracks Weekly and Daily Symptom Variations during Deep Brain Stimulation Therapy for Depression. IEEE J. Biomed. Health Inform. 1–14 (2025) doi:10.1109/JBHI.2025.3566057.

114. Sellers, K. Towards individualized deep brain stimulation: A stereoencephalography-based workflow for unbiased neurostimulation target identification. bioRxiv (2025).

115. Mehrabian, A. Pleasure-arousal-dominance: A general framework for describing and measuring individual differences in Temperament. Curr. Psychol. 14, 261–292 (1996).

116. Fontaine, J. R. J., Scherer, K. R., Roesch, E. B. & Ellsworth, P. C. The World of Emotions is not Two-Dimensional. Psychol. Sci. 18, 1050–1057 (2007).

117. Qasim, S. E., Fried, I. & Jacobs, J. Phase precession in the human hippocampus and entorhinal cortex. Cell 184, 3242–3255.e10 (2021).

118. Geva-Sagiv, M. et al. Augmenting hippocampal–prefrontal neuronal synchrony during sleep enhances memory consolidation in humans. Nat. Neurosci. 26, 1100–1110 (2023).

119. Fischl, B. FreeSurfer. NeuroImage 62, 774–781 (2012).

120. Tadel, F., Baillet, S., Mosher, J. C., Pantazis, D. & Leahy, R. M. Brainstorm: A User-Friendly Application for MEG/EEG Analysis. Comput. Intell. Neurosci. 2011, 1–13 (2011).

121. The Qt Company. Qt6. (2022).

122. Kleiner, M. What’s new in Psychtoolbox-3? (2007).

123. Lang, P.J., Bradley, M.M., & Cuthbert, B.N. International affective picture system (IAPS): Technical manual and affective ratings. NIMH Cent. Study Emot. Atten. 3, 39–58 (1997).

124. Dan-Glauser, E. S. & Scherer, K. R. The Geneva affective picture database (GAPED): a new 730-picture database focusing on valence and normative significance. Behav. Res. Methods 43, 468–477 (2011).

125. Pengfei, X., Yuxia, H. & Yuejia, L. Establishment and assessment of native Chinese affective video system. Chin. Ment. Health J. 24, (2010).

126. Chen, W., Wang, Y. & Yang, Y. Efficient Estimation of Directed Connectivity in Nonlinear and Nonstationary Spiking Neuron Networks. IEEE Trans. Biomed. Eng. 71, 841–854 (2024).

127. Ian Goodfellow, Yoshua Bengio, & Aaron Courville. Deep Learning. (MIT Press, 2016).

128. Cawley, G. C. & Talbot, N. L. C. On Over-fitting in Model Selection and Subsequent Selection Bias in Performance Evaluation. The Journal of Machine Learning Research 2079–2107 (2010).

129. Maaten, L. & Hinton, G. Visualizing Data using t-SNE. J. Mach. Learn. Res. 9, 2579–2605 (2008).

130. Maulik, U. & Bandyopadhyay, S. Performance evaluation of some clustering algorithms and validity indices. IEEE Trans. Pattern Anal. Mach. Intell. 24, 1650–1654 (2002).

131. Cohen, M. X. Analyzing Neural Time Series Data: Theory and Practice. (The MIT Press, Cambridge, Massachusetts, 2014).

132. FFmpeg Developers. FFmpeg. (2024).

133. Introduction to Meta-Analysis. (Wiley, Chichester, 2013).

